# De novo genome assembly of four Andean potato weevil species (*Premnotrypes, Rhigopsidius*), the primary agricultural pest of the potato in South America

**DOI:** 10.1101/2023.12.13.571405

**Authors:** Kelsey C. Jorgensen, Obed A. Garcia, Jesús Alcázar, Kimberly K.O. Walden, Abigail W. Bigham, Norma Mujica Morón, Clorinda Vergara Cobián, Julie J. Lesnik, Chuanzhu Fan

## Abstract

The Andean potato weevil complex are the most widespread and serious insect pests to potato crops in the Andes. More broadly, genomic assemblies of insect pests are currently lacking in agricultural research, especially those from the order Coleoptera. These genome data are essential for identifying potential underlying mechanisms important to biological control strategies and food security in the highlands. Here, we present the *de novo* genome assemblies for four species of the Andean potato weevil complex: *Premnotrypes vorax, P. suturicallus*, *P. latithorax*, and *Rhigopsidius piercei*. Genome assemblies exceeded the average size of those from the order Coleoptera and were highly repetitive: for *P. vorax* (1.33 Gb, 71.51% repetitive), *P. latithorax* (623 Mb, 59.03% repetitive), *P. suturicallus* (1.23 Gb, 70.19% repetitive), and *R. piercei* (1.55 Gb, 71.91% repetitive). We examined genomic regions related to metabolic potato plant detoxification and insecticide resistance using the available Colorado potato beetle (*Leptinotarsa decemlineata*) genome annotations as a guide. Our analysis of these weevil genomes identified chemosensory receptors and odorant binding proteins that could be related to detecting their hosts, the potato plant (*Solanum tuberosum*), as well as many genomic regions involved in subverting pesticide resistance. We have generated the first whole-genome assemblies of the Andean potato weevil complex that will be foundational for future agricultural pest management and entomological research in South America.

**Author Summary:** Within the South American Andean mountains the Andean potato weevil insects are the most widespread and serious pests to potatoes, destroying around 89% of potato harvests a year when insecticides are not used. Here, we collected and performed whole-genome sequencing for the first time for four Andean potato weevil species: *Premnotrypes vorax, P. suturicallus*, *P. latithorax*, and *Rhigopsidius piercei*. After analysis of these genome assemblies, we found that they were large and highly repetitive compared to other published beetle genome data in the order Coleoptera. After further examination of these genome assemblies, we found regions related to metabolic potato plant detoxification, insecticide resistance, and chemosensory and odorant binding protein receptors that could be related to detecting potato plants. These genomic identifications provide novel molecular insight into regions associated with insecticide resistance, metabolic abilities, and environmental receptors, and can serve as a future valuable resource in classifying phylogenetic relationships as well as identifying regions of interest for improved pest management for potato farmers.

## Introduction

The Andean potato weevil complex is the most prevalent potato pest in the South American Andes (1–3), destroying an average of 89% of potato tuber harvests without pest control measures (4). This complex has a unique speciation history limited to where the potato was first domesticated in the Andean highlands (5–7), and consists of 14 species across three genera (*Premnotrypes*, *Rhigopsidius*, *Phyrdenus*), although inclusion of three of these species are disputed. Andean potato weevil genera belong to the subfamily Entiminae (∼12,220 species), within the large family Curculionidae (∼83,000 species), and order Coleoptera (∼400,000 species) (8–10). Of the confirmed Andean potato weevil species, ten are found in the Peruvian Andes, one species only in Bolivia, and four species ranging more broadly from Argentina to Venezuela (**Table 1**). The most widespread pest species within the Andean potato weevil complex are *P. latithorax* (Pierce 1914), *P. suturicallus* (Kuschel 1956), and *P. vorax* (Hustache 1933). It is estimated that these three species cause an average of 30% to 50% of the total crop yield loss and are most prolific in Peru (2, 11, 12). Species in this complex are cold adapted with optimal fecundity between 11°C and 15°C and severely reduced fecundity in temperatures above 20°C. At temperatures above 25°C, Andean potato weevils are unable to survive. Furthermore, all species in this complex except one disputed species cannot fly. Together, these constraints restricts their habitats to elevations between about 2,100 to 4,700 meters (1, 11).

**Table 1.** Andean potato weevil Species and Geographic Distributions.

| Species | Geographic Region from Literature | Elevation range (meters) |
| --- | --- | --- |
| *Premnotrypes vorax* | Northern Peru, Ecuador, Western Columbia, Western Venezuela | 3135-3519 |
| *P. solanivorax* | Northern Peru | 3390-3397 |
| *P. solani* | Central Peru | 3069-3357 |
| *P. fractirostris* | Central Peru | 3263-3713 |
| *P. suturicallus* | Central Peru | 3296-3560 |
| *P. piercei* | Central Peru | 3302-3263 |
| *P. pusillus* | Central Peru, Southern Peru | 4090-4208 |
| *P. solaniperda* | Southern Peru, Northern Bolivia | 3230-3834 |
| *P. latithorax* | Southern Peru, Northern Bolivia, Chile | 3230-4183 |
| *P. zischkai* | Bolivia | 3900-4000 |
| *Rhigopsidius piercei* | Southern Peru, Bolivia, Northern Chile, Northern Argentina | 3783-3834 |
| Disputed Species | Geographic Region from Literature | Elevation Range (meters) |
| *P. sanfordi* | Peru | Unknown |
| *P. clivosus* | Bolivia | Unknown |
| *Phyrdenus muriceus* | Argentina, Bolivia | 2100-2400 |

Species of the Andean potato weevil complex depend completely upon the potato plant (*Solanum tuberosum*) to complete their life cycle (**Figure 1**). Adult females lay their eggs in the soil close to the potato plant, so that the neonate larvae can burrow into the nearby potato tubers after hatching. Larvae feed on tubers for about 40 days, after which they abandon the potato and pupate deep (about 30cm) into the nearby soil. Emergence of the adult weevil from the soil is stimulated during the rainy season, generally between October through January. The exception to this behavior is *Rhigopsidius piercei* (Heller 1906), which completes its pupation inside the tuber and emerges as an adult weevil (1). Overall, their life cycle is synchronized over the course of one year and culminates with the potato harvest period. Adult Andean potato weevils will remain within the same potato field if plants are available (1, 2, 4). Otherwise, they will move to adjacent potato fields. They can migrate by walking distances up to 500 meters but are unable to cross running bodies of water (11, 13, 14).

**Figure 1.**
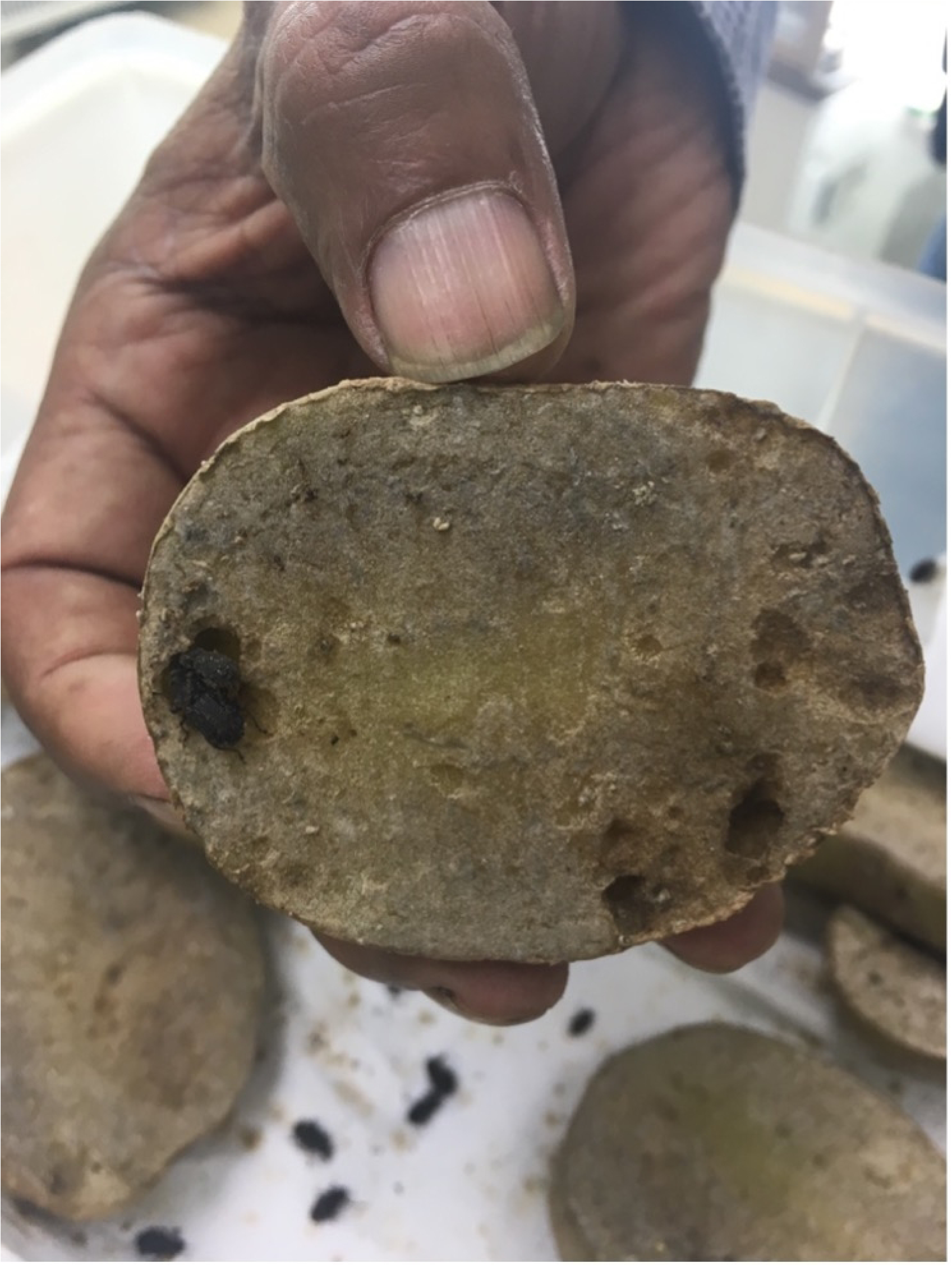
Andean potato weevils inside potato. Photo taken by K. Jorgensen shows Andean potato weevils feeding within a potato and the characteristic circular divots of previous feeding locations.

The primary form of Andean potato weevil pest control is the application of highly toxic pesticides that have detrimental health and environmental effects (4, 15–17). Non-pesticide forms of control also exist including the construction of plastic barriers (>25cm) around fields, crop rotation, and increased distances between potato fields (2, 18, 19). Furthermore, both pesticide and non-pesticide forms of control range in their efficacy. Andean potato weevil species can still destroy up to 70% of tuber crops when insecticides are applied, around 16.6%- 44.7% of crops when plastic barriers are used, and around 9.4%-81.1% of crops when plastic barriers are used in conjunction with insecticides (2, 12, 20).

Other efforts, such as the development of a pest-resistant potato cultivar or increasing the number of natural predators, have also been unsuccessful (2, 19, 21). For the latter, this is partly due to the lack of specific Andean potato weevil pest complex predators. Various carabid beetles (*Harpalus turmalinus*, *Notiobia schnusei*, *Harpalus turmalinus*), or toads, birds, and spiders will opportunistically consume their eggs and larvae, and the weevils are also susceptible to entomopathogenic fungi (*Beauveria bassiana*, *Beauveria brongniartii*) and nematode parasites (*Heterorhabditis* sp. and *Steinernema* sp.) (2, 11, 22). Some of the most successful pest control strategies are instead emerging through studying and manipulating the genetic diversity that enables weevils to subvert pesticides or metabolize toxic compounds present in potato plants.

A genomic characterization of the Andean potato weevil complex would be of great benefit to pest control and agricultural management. Pest analyses of other insect genomes suggest the mechanisms of underlying metabolic interdependencies between co-evolutionary organisms could represent potential targets for agricultural pest management (23). For example, metagenomic analysis found that the long-horned passalid beetle (*Odontotaenius disjunctus* Illiger 1800) had a uniquely structured microbiome that allowed the insect to consume nutrient-poor wood (24). As such, manipulating the insect gut microbiome by leveraging plant-mediated expression of antimicrobial peptides is an alternative strategy to decreasing insect fitness (25). Other potential targets for insect pest management are the development of bioinsecticides, or the application of naturally occurring biological substances to repel pests. When considering the monophagous nature of the Andean potato weevil complex, developing a bioinsecticide solution would be a highly beneficial alternative to the toxic insecticides currently used by Andean farmers (4, 15–17). In this vein, the most successful pathogen used for insect control is the bacterium *Bacillus thuringiensis* (Bt), which kills the insect in its larval stage by disrupting the midgut tissue and invoking septicemia. Bt proteins produce insecticidal pore-forming proteins called Cry or Cyt toxins that can be activated in insects that consume them after insecticide application, or more commonly, introduced into transgenic crops (15). In one successful case, transgenic cotton producing the *Bt* toxin Cry1Ac killed almost 100% of the destructive cotton pink bollworm larvae (*Pectinophora gossypiella* Saunders 1844) pest in the southwestern United States (26). The opportunity to explore these underlying metabolic mechanisms in insect pests is only possible with the increased data and coverage generated by whole-genome sequencing (23).

While the behavior and morphology of Andean potato weevil species have been well-characterized, very few of the species have been sequenced. Partial mitochondrial and nuclear rRNA gene sequences (*18S*, *28S*, *16S*, *COI*) for *P. latithorax* and *R. piercei* were published as part of a larger phylogeny of Entiminae weevils (27). A transcriptome assembly for *P. vorax* is publicly available as unassembled reads (28). About 1,000 species in the Andean potato weevil genera subfamily Entiminae have been sequenced to varying levels, including some mitogenomes, and made publicly available on NCBI (29–33). The genome sizes in this highly diverse order can range from 160 to 5,020 Mb, with an average of 760 Mb (34). Here, four *de novo* genomes from the Andean potato weevil complex, including three of its most widespread pest species, were sequenced and assembled for the benefit of future agricultural, pest, and entomological research. To date, this is the first study to feature a whole-genome assembly of any species of the Andean potato weevil complex.

## Results

### Genome Assembly and Quality Assessment

Illumina sequencing yielded the following read-pair counts from each sample: *P. vorax* – 209,818,864, *P. latithorax* – 238,387,163, *P. suturicallus* – 210,931,219, and *R. piercei* – 252,278,491. Three of the genome assembly sums were larger than predictions by GenomeScope 2.0 (35), and larger than the coleopteran average of 760 Mb (34).

GenomeScope 2.0 outputs predicted sizes of 619 Mb for *P. vorax* (53D), 711 Mb for *P. latithorax* (69J), and 613 Mb for *P. suturicallus* (28J). Only *R. piercei* (90H) was predicted to be larger than the average size for the Coleoptera order at 957 Mb (**Figure S1**). The final assemblies summed to 1.33Gb for *P. vorax*, 1.23 Gb for *P. suturicallus*, 1.55 Gb for *R. piercei*, and a smaller 623 Mb for *P. latithorax*.

For *R. piercei* (90H), there were over 350,000 scaffolds with a genome size of 1.55 Gb and scaffold N50 around 6.4 kb. *P. vorax* (53D) had over 470,000 scaffolds with a genome size of 1.33 Gb, and scaffold N50 around 3.4 kb. *P. latithorax* (69J) had a genome size of 623 Mb, over 300,000 scaffolds, and a scaffold N50 of 2.2 kb. The final *P. suturicallus* has a genome size of 1.23 Gb, over 350,000 scaffolds, and scaffold N50 of 4.8 kb. BUSCO completeness scores ranged from 39.6% for *R. piercei*, 54.2% for *P. vorax*, 45.9% for *P. latithorax*, and 52.7% for *P. suturicallus* (**Table 2**).

**Table 2.**
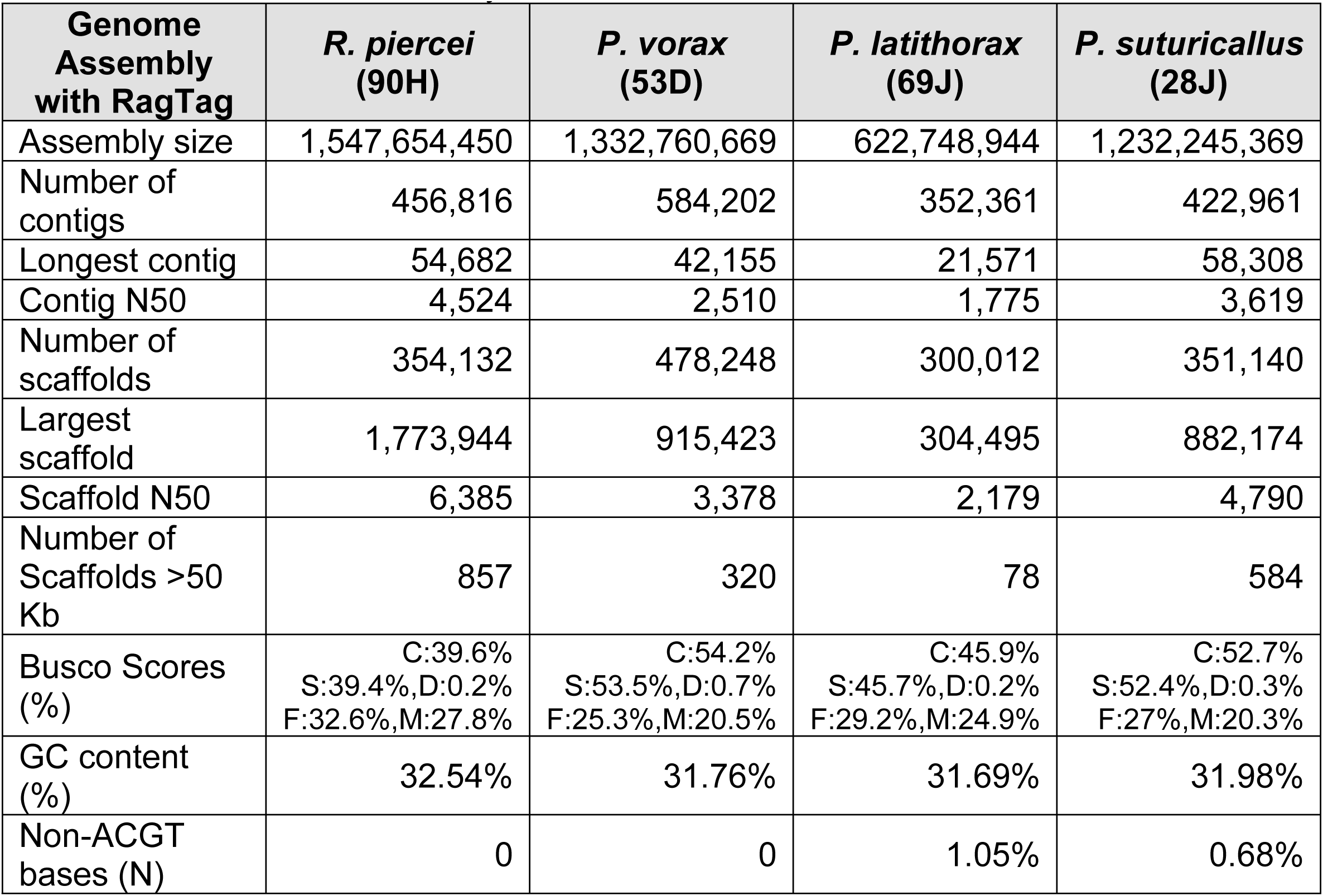
Final Genome Assembly Metrics.

| Genome Assembly with RagTag | *R. piercei* (90H) | *P. vorax* (53D) | *P. latithorax* (69J) | *P. suturicallus* (28J) |
| --- | --- | --- | --- | --- |
| Assembly size | 1,547,654,450 | 1,332,760,669 | 622,748,944 | 1,232,245,369 |
| Number of contigs | 456,816 | 584,202 | 352,361 | 422,961 |
| Longest contig | 54,682 | 42,155 | 21,571 | 58,308 |
| Contig N50 | 4,524 | 2,510 | 1,775 | 3,619 |
| Number of scaffolds | 354,132 | 478,248 | 300,012 | 351,140 |
| Largest scaffold | 1,773,944 | 915,423 | 304,495 | 882,174 |
| Scaffold N50 | 6,385 | 3,378 | 2,179 | 4,790 |
| Number of Scaffolds >50 Kb | 857 | 320 | 78 | 584 |
| Busco Scores (%) | C:39.6%<br>S:39.4%,D:0.2%<br>F:32.6%,M:27.8% | C:54.2%<br>S:53.5%,D:0.7%<br>F:25.3%,M:20.5% | C:45.9%<br>S:45.7%,D:0.2%<br>F:29.2%,M:24.9% | C:52.7%<br>S:52.4%,D:0.3%<br>F:27%,M:20.3% |
| GC content (%) | 32.54% | 31.76% | 31.69% | 31.98% |
| Non-ACGT bases (N) | 0 | 0 | 1.05% | 0.68% |

Kraken2 analysis suggested that contaminant reads were not highly abundant in the raw sequence data for the four species (36). For *R. piercei*, 97.69% were labeled as ‘unclassified’ or potential hits to the target organism, and 2.31% of sequences were assigned to bacteria or virus, plant, or other/unknown. For *P.* vorax, 98.75% of sequences were identified as the target organism and 1.25% of sequences classified as contaminants. Both *P. suturicallus* and *P. latithorax* libraries had higher percentages of contaminant reads removed. *P. suturicallus* had 93.67% reads and *P. latithorax* had 95.44% reads identified as the target organism, with 6.33% and 4.56% reads identified as contaminants, respectively.

After sensitive-MEGAHIT assembly, BlobTools was used to identify any further contaminants in the weevil assemblies. For *R. piercei*, 95.28% out of 96.75% contigs were classified as ‘no hit’ and Arthropoda (**Table S1**; **Figure S2**). The categories ‘no hit’ and any general Arthropoda sequences are considered the de novo target organism since they do not share a significant match in the UniProt database. A total of 0.15% of sequences were classified as either bacterial or parasitic in origin, with less than 0.08% categorized as potential fungal contaminants. The remaining 1.25% were classified as possible plant sequences or ‘other’ unknown contaminants. However, hits to plant sequences are often spurious and could be derived from insect ribosomal DNA sequences. These sequences were isolated and examined more closely, and true hits to plant sequences were removed along with hits to contaminant bacterial and viral sequences. Similar levels were observed for the other three species. For the *P. suturicallus*, 97.87% out of 99.06% contigs were classified as Arthropoda and ‘no hit’, with 1.2% as contaminants. For *P. latithorax*, 96.04% out of 96.78% contigs were classified as ‘no hit’ and Arthropoda, with only 0.73% as contaminants. Lastly for *P. vorax*, 95.11% out of 96.77% contigs were classified as ‘no hit’ and Arthropoda, with 1.66% as possible contaminants. Results using Dedupe with BBTools indicated there were no duplicated sequences that needed to be removed from any of the four assemblies (**Table S2**).

After scaffolding with SOAPdenovo-fusion, the total number of scaffolds/contigs decreased while the other metrics remained stable (**Tables S2-3**). The longest scaffold increased on average by 9,314 bp, and BUSCO ortholog prediction scores slightly improved for all four assemblies. Further scaffolding with L_RNA_Scaffolder using the *P. vorax* transcript open reading frames as guides resulted in significant improvement for all four assemblies (**Tables S3-4**). The longest scaffold increased on average by 37,946 bp, and the number of scaffolds >50 kb rose from around 3 to 44. Complete BUSCOs improved by 14.2% on average, and fragmented BUSCOs decreased by 6.18%.

For the last round of scaffolding in RagTag, four independent Curculionid genome assemblies and one merged assembly were used in reference-guided scaffolding with both *R. piercei* and *P. vorax*. Independent use of the *Listronotus bonariensis* (Argentine stem weevil Kuschel 1955) reference produced the best metric improvements for both species (**Table S5**). After reference-guided scaffolding was complete for all species, BUSCO scores improved for all four species (**Table 2**). The length of the longest scaffold increased on average by 879,304 bp, complete BUSCOs improved by about 1.4%, and the number of scaffolds >50 kb rose drastically from around 44 to 460 for the four weevil assemblies.

### Repeat Elements

Insect genomes are prone to having highly repetitive sequences, which can lead to more challenging genome assembly (37). Transposable elements (TE) are one type of highly repetitive DNA sequence believed to have considerable influence on the evolution of the genome. High TE content can be a hurdle to genome assembly due to their high copy numbers and complex nesting structures produced by new TE insertions (38). Repetitive content was estimated at 59.03% with *P. latithorax,* to 71.91% with *R. piercei*, 71.51% with *P. vorax*, and 70.19% with *P. suturicallus* (**Table 3**). From the 100 longest scaffolds of the *R. piercei* assembly, 63.28% were classified as transposable elements using RepeatMasker. Two of the other Andean potato weevil assemblies also had the majority of their longest 100 scaffolds match TEs, with 55.48% from *P. vorax* and 58.31% from *P. suturicallus*. Just under half (41.85%) of the 100 longest scaffolds from the *P. latithorax* assembly matched TEs. TE content is also positively correlated to genome assembly size (39), and it is possible that these weevil genome assemblies could contain even higher amounts of repetitive regions if they were analyzed in the future with long-read sequencing and improved scaffolding technologies.

**Table 3.** Repeat Elements of Final Genome Assemblies.

| ***R. piercei* (90H)**<br>**Repeat Types** | **Number of**<br>**Elements** | **Length**<br>**occupied (bp)** | **Percentage of**<br>**Sequence** |
| --- | --- | --- | --- |
| SINE | 15,424 | 2,454,676 | 0.16% |
| LINE | 802,792 | 26,447,977 | 17.09% |
| LTR | 52,271 | 28,660,165 | 1.85% |
| DNA transposons | 561,205 | 156,494,454 | 10.11% |
| Unclassified | 2,762,295 | 633,734,852 | 40.95% |
| Small RNA | 22,332 | 4,053,112 | 0.26% |
| Total bases masked | 4,216,319 | 1,112,979,060 | 71.91% |
| ***P. vorax* (53D)**<br>**Repeat Types** | **Number of**<br>**Elements** | **Length**<br>**occupied (bp)** | **Percentage of**<br>**Sequence** |
| SINE | 6,769 | 1,075,500 | 0.08% |
| LINE | 832,140 | 244,081,151 | 18.31% |
| LTR | 36,763 | 11,091,824 | 0.83% |
| DNA transposons | 890,028 | 276,514,979 | 20.75% |
| Unclassified | 2,192,505 | 412,135,273 | 30.92% |
| Small RNA | 897 | 91,446 | 0.01% |
| Total bases masked | 3,959,102 | 953,090,590 | 71.51% |
| ***P. latithorax* (69J)**<br>**Repeat Types** | **Number of**<br>**Elements** | **Length**<br>**occupied (bp)** | **Percentage of**<br>**Sequence** |
| SINE | 11,331 | 1,663,233 | 0.27% |
| LINE | 348,156 | 82,312,970 | 13.22% |
| LTR | 17,813 | 5,589,527 | 0.9% |
| DNA transposons | 379,440 | 86,793,027 | 13.94% |
| Unclassified | 1,265,567 | 185,798,544 | 29.84% |
| Small RNA | 13,009 | 2,354,730 | 0.38% |
| Total bases masked | 2,035,316 | 367,619,230 | 59.03% |
| ***P. suturicallus* (28J)**<br>**Repeat Types** | **Number of**<br>**Elements** | **Length**<br>**occupied (bp)** | **Percentage of**<br>**Sequence** |
| SINE | 3,438 | 483,858 | 0.04% |
| LINE | 571,549 | 175,543,344 | 14.25% |
| LTR | 42,808 | 16,213,230 | 1.32% |
| DNA transposons | 766,722 | 228,616,991 | 18.55% |
| Unclassified | 2,590,240 | 437,097,018 | 35.47% |
| Small RNA | 3,404 | 352,912 | 0.03% |
| Total bases masked | 3,978,161 | 864,897,315 | 70.19% |

### Environmental receptors and digestive functions

All species of the Andean potato weevil complex are monophagous and use environmental signals to sense the *Solanum* potato plant. These environmental signals can include chemical, olfactory, visual, or tactile cues, which are encoded by olfactory, light sensory, and neural receptor genes (34, 40–44). Here, loci previously associated with these environmental cues and labeled as either protein-coding genes or predicted protein-coding genes in the available Colorado potato beetle (*Leptinotarsa decemlineata* Say 1824) genome assembly (NCBI PRJNA854273) were used to identify similar gene regions in our four Andean potato weevil species using discontiguous MEGABLAST (34, 45) (**Tables S6-10**).

Odorant receptors are believed to be the dominant way for insects to detect olfactory signals, and the heterodimeric channels are formed by an odorant-specific receptor protein (OR) and the olfactory receptor co-receptor (Orco) (42). ORs evolve rapidly with low levels of homology between insect families and Orcos are critical for insect olfactory transduction.

Gustatory receptor genes (GRs) encode seven-transmembrane proteins and play a primary part in an insect’s bitter receptors and ability to detect insecticides such as DEET (N.N-diethyl-meta-toluamide) (46). Ionotropic receptors (IRs) are highly conserved and similar to GRs as they are involved in both olfaction and gustation (42). One group of ionotropic receptors, e.g., Ir21a, 40a, 68a, and 93a, are particularly associated with sensing temperature (34). Odorant binding proteins (OBPs) are similar to ORs, GRs, and IRs in insects as they are part of chemosensory gene families that contribute to oligophagous insects’ detection of organic plant compounds through olfactory perception (42). These families are large in insect genomes and can often consist of hundreds of genes (34). We identified an average of 32 GR gene regions, 39 ORs, 20 IRs, and 30 OBPs across the four Andean potato weevil genomes, fewer than those annotated for *L. decemlineata* (65 GRs, 45 ORs, 32 IRs, and 41 OBPs) (**Table S6**).

The opsin gene family is part of the G-protein-coupled transmembrane receptors and expressed in the retina of insects (47, 48). These photoreceptors are associated with detecting yellow light and enable *L. decemlineata’s* ability to sense the yellow flowers of its host plant *S. rostratum* (34). There was an average of 29 opsins identified across the four Andean potato weevil genomes related to UV-sensitivity.

Carbohydrate active enzymes (CAZy) are involved with breaking down complex carbohydrates and categorized into five major classes with hundreds of enzymes (49). To narrow the search within these hundreds of genes, any region related to three particular gene families (GH28, GH45, GH48) were identified in the four Andean potato weevil assemblies.

GH28, GH45, and GH48 are CAZy gene families linked to plant cell wall carbohydrate digestion since they contain enzymes that break down leaf pectin and cellulose (50). There was an average of five GH28, one GH45, and one GH48 CAZy gene families identified in the *R. piercei*, *P. vorax*, *P. latithorax*, and *P. suturicallus* genomes, respectively (**Table S6**).

Cysteine proteases are part of *L. decemlineata’s* ability to digest toxic potatoes and may even have contributed to a broader coleopteran ability to expand digestive functions beyond initial host plants (34). Both CAZy enzymes and cysteine peptidases are greatly diverse digestive enzymes that show high expression in beetle gut tissue and play a major role in phytophagy (34, 50). There were about 39 cysteine peptidase genes on average across the four Andean potato weevil genomes (**Table S6**).

Cytochrome P450 monooxygenases (CYPs) genes, particularly CYP6 and CYP12, are associated with both insecticide resistance and detoxification of plant allelochemicals in *L. decemlineata* and other insects (34). There was an average of 103 CYP genes across the four weevil genomes, which are less compared to the beetle *L. decemlineata* (145 genes) (34, 45) (**Table S6**).

### Insecticide Resistance

Insects that feed on potato plants must develop complex digestive abilities since solanaceous plants contain highly toxic compounds such as steroidal alkaloids and glycoalkaloids (51–53). The physiological mechanisms involved in detoxifying potato plant compounds could also be involved in subverting pesticide resistance (53, 54). Target regions involved in both metabolic potato plant detoxification and insecticide resistance were annotated in all four Andean potato weevil assemblies and listed in **Tables S6-10** (34, 55, 56).

Carboxylesterase (CarE) genes are a class of detoxification genes that may play a role in insecticide resistance by catalyzing xenobiotic and endogenous compounds. CarE enzymes are also present in both bee and wasp venom and could be allergen proteins (57). Glutathione S-transferases (GSTs) are one of the main detoxification enzyme systems in insects, including *L. decemlineata*, and have been shown to resist insecticides through direct metabolism in fruit flies (34, 58). Examination of these potential insecticide resistance regions found about 34 CarE enzymes and 26 GSTs across the four Andean potato weevil genomes (**Table S6**).

Other targets for the development of insecticides focus more specifically on manipulating environmental sensors. The transient receptor potential channels (TRPs) are proteins involved in the full range of environmental sensors in insects including hearing, smell, taste, sight, temperature, and pheromones. Around 13 to 14 TRP genes are targeted by common commercial insecticides and found in insect stretch receptor cells. Histamine-gated chloride channels (HisCls) are the transmitter of arthropod photoreceptors and thoracic mechanoreceptors (59), and the cation-gated nicotinic acetylcholine receptors (nAChRs) are expressed in the central nervous system of insects (60). In the four Andean potato weevil genomes, there is a similar number of these receptors present including 17 TRPs, 1 HisCL, and 19 nAChRs compared to *L. decemlineata* (19 TRPs, 2 HisCLs, 20 nAChRs) (**Table S6**). The use of insecticides in agriculture has inevitably led to insecticide resistance, and the γ-amino butyric acid gated anion (GABA) channels is one region that has been associated with insecticide resistance to dieldrin (*RDL*), and other organic insecticides such as cyclodienes in flies (34). Cuticle proteins have also been shown to have imidacloprid resistant traits following RNAi knockdown experiments in *L. decemlineata* (61). There were similar numbers of GABA receptors (average 15) and fewer cuticle genes (average 61) identified in the four Andean potato weevil genomes compared to *L. decemlineata* (17 GABAs, 90 cuticle genes) (34, 45, 62) (**Table S6**).

### Synteny Analysis

Whole-genome synteny analysis was performed using protein reading frames in MUMmer4 for each of the Andean potato weevil genomes and the chromosome-level published genome of *L. decemlineata*. We extracted the olfactory, light sensory, neural receptor, and insecticide resistance gene regions previously identified using discontiguous MEGABLAST from this whole genome synteny to determine the degree of gene order conservation between each Andean potato weevil genome and the corresponding region within the *L. decemlineata* genome. These gene coordinates were plotted using CIRCOS after being classified into the following classes: CarE, CAZy, CYPs, cuticles, cysteine, GABAs, GPCRs, GRs, GSTs, HisCLs, IRs, nAChRs, OBPs, ORs, and TRPs (**Figure 2**). Although detectable synteny tends to decrease with more fragmented assemblies, we still were able to detect a high degree of synteny in our results (63).

**Figure 2.**
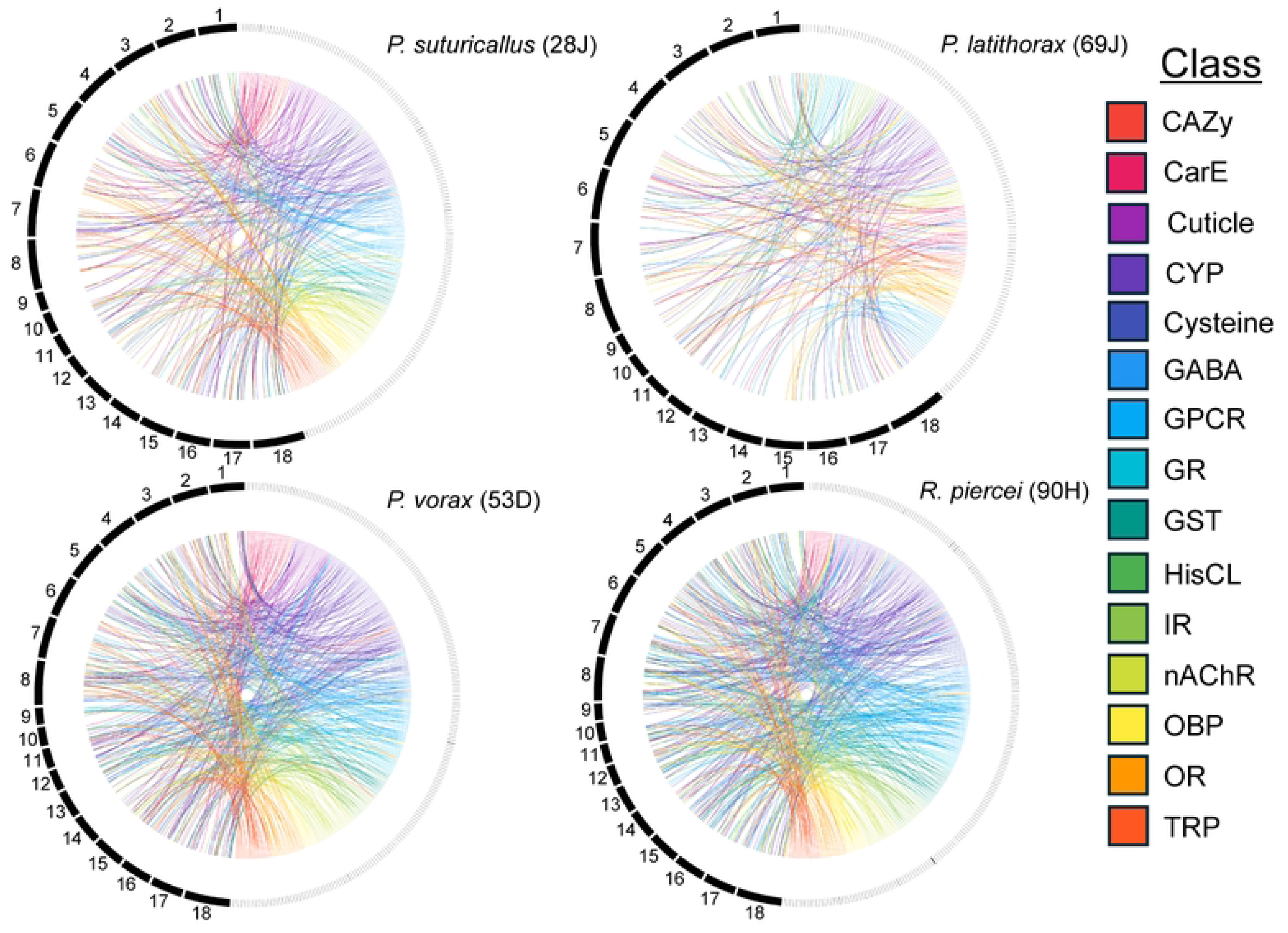
Synteny analysis between Andean potato weevil genomes and *L. decemlineata.* For each plot, heavy black lines indicate the *L. decemlineata* chromosome level assembly, while the smaller marks indicate Andean potato weevil contigs. There is a large amount of synteny between *L. decemlineata* and *R. piercei* and *P. vorax*, and the least amount of shared synteny with *P. latithorax*. However, our synteny analysis would have been affected by final assembly metrics, of which *P. latithorax* had the smallest contig and scaffold N50s.

High sequence synteny is indicated through the high number of links between regions, here GABAs, nAChRs, and TRPs share particularly high syntetic clusters with the same region of *L. decemlineata*. The CAZy gene families showed high synteny between *P. suturicallus* and *L. decemlineata*, very little for *P. vorax* and *L. decemlineata*, and none for both *R. piercei* and *P. latithorax* with *L. decemlineata*. For HisCLs, *R. piercei* shared synteny with *L. decemlineata*, whereas three of the four Andean potato weevil species (*P. suturicallus*, *P. vorax*, and *P. latithorax*) shared none.

Both *R. piercei* and *P. vorax* shared higher amounts of syntetic links with *L. decemlineata* for cysteine peptidase genes, G-protein-coupled transmembrane receptors, and IRs, while *P. suturicallus* and *P. latithorax* shared lower amounts of synteny with *L. decemlineata* for these queries. For GSTs and OBPs, *R. piercei* shared higher amounts of syntetic links with *L. decemlineata* compared to the other three weevil species. The remaining CarE, cuticle, CYPs, GRs, and OR genes all share moderate synteny across all four Andean potato weevil genomes and *L. decemlineata*.

While the Andean potato weevil genomes are not annotated to the chromosome-level, the high sequence synteny between the regions indicated and *L. decemlineata* suggests that they ultimately share similar gene functions. Additionally, any occurrence of highly shared synteny signifies an elevated degree of evolutionary protein conservation, since these four Andean potato weevil species and *L. decemlineata* are not closely related within the order coleopteran.

## Discussion

Species of the Andean potato weevil complex are the most severe pests of the potato in the Andes, where infestations destroy up to 89% of potato harvests when insecticides are not applied. There is an urgent need for an effective pest control strategy, since up to 70% of tubers are still infested even after Andean farmers apply highly toxic pesticides (2, 4). Three of the four Andean potato weevil genome assemblies presented here, *P. vorax*, *P. latithorax*, and *P. suturicallus*, are the most widespread pests of this complex and can cause around 30% to 50% of total potato crop loss (2, 11, 12). An emerging focus in pest control is the development of a bioinsecticide specific to the insect species or complex when other forms of pest control are ineffective. Here, we present the first *de novo* genome assemblies of the Andean potato weevil complex with the species *Premnotrypes vorax, P. latithorax, P. suturicallus* and *Rhigopsidius piercei*. The genome assemblies for three of these four weevils are exceptionally large for Coleoptera at 1.55 Gb (*R. piercei*: x12-fold read coverage), 1.33 Gb (*P. vorax*: x12-fold read coverage), and 1.23 Gb (*P. suturicallus*: x13-fold read coverage). *P. latithorax* is an average genome size for coleopterans at 623 Mb (x29-fold read coverage). Initial annotation analysis of these four weevil assemblies provided novel molecular insight into genomic regions associated with insecticide resistance or environmental receptors and can serve as a future valuable resource in identifying regions of interest for improved pest management.

The lower BUSCO completeness scores for these weevil genomes are likely the result of short-read sequencing libraries and the absence of additional long-read linkage information (64). The draft assembly of the agricultural pest Asiatic rice borer moth (*Chilo suppressalis* Walker 1863) had similar statistics, as it was generated using Illumina HiSeq 2000 and had a final size of 824 Mb with scaffold N50 of around 5.2 kb. Despite these limitations, annotation of the *C. suppressalis* genome featured many chemosensory and odorant binding proteins as well as genes along metabolic pathways. Despite the lower BUSCO completeness scores of these four Andean potato weevil assemblies, early annotation has also identified many gene regions related to chemical, olfactory, visual, or tactile cues, or involved in subverting pesticide resistance.

Genome profiling has also been widely used for the classification of insect species and created new insights into their evolutionary relationships. For the Andean potato weevil complex, the inclusion of three weevil species is currently disputed. Two species (*P. clivosus* and *P. sanfordi*) were described from single deceased specimens (65). It is possible that *P. sanfordi* could be reclassified as part of *P. latithorax* based on its morphological description. Importantly, both *P. clivosus* and *P. sanfordi* have never been observed in the field since these initial descriptions and could even represent morphological variation of a previously described species. The third disputed species, *Phyrdenus muriceus,* has vastly different morphological and behavioral characteristics from the rest of the complex. *P. muriceus* can fly, is adapted to live at lower altitudes (2,100 to 2,400 meters) and has a geographic range across both North and South America. *P. muriceus* is not dependent on the potato for the completion of its life cycle and can feed on other crops such as tomatoes and eggplants. For these reasons, scholars have argued that *P. muriceus* belongs in a classification separate from the Andean potato weevil complex (1, 11, 65). The genomic data generated here is foundational for future studies classifying the phylogeny of the Andean potato weevil complex and the relatedness between disputed and confirmed species.

## Limitations

There are several factors complicating improved annotation of these assemblies. First, three of the four weevil genomes were larger than the average size for the order Coleoptera (760 Mb), ranging from 1.23 Gb to 1.55 Gb (34). For context, the initial genome assembly of the highly studied Colorado potato beetle (*L. decemlineata*) was 1.17 Gb (34), and a more recent chromosome-level assembly is about 1.01 Gb (45). Underlying high levels of heterozygosity (1% or above) can also result in allelic differences resembling paralogy in fragmented assemblies even with high sequencing quality (66). The high heterozygosity predicted by GenomeScope for each of these species (*R. piercei*: 7.56%, *P. vorax*: 8.02%, *P. latithorax*: 7.68%, *P. suturicallus*: 7.62%) additionally contributed to difficult genome assembly with the result of lower BUSCO completeness scores. While the diamondback moth (*Plutella xylostella*) genome is only 343 Mb, its high degree of heterozygosity made successful genome assembly impossible until You, Yue (67) implemented a later fosmid-to-fosmid sequencing strategy combined with whole-genome sequencing.

Challenges over securing international permits and collection also led to downstream complications with assembly. Using inbred insects is ideal for genome assembly as it can significantly reduce heterozygosity and allelic diversity. However, all these species were wild caught since the only breeding program for Andean potato weevils was discontinued in 2017 at the Centro Internacional de la Papa (CIP) in Peru. In addition, pupae are often the best life-stage to isolate DNA from insects as they do not contain the chitinous material (exoskeletons) of adults or large quantities of gut microbiota (ingested food) that can contaminate assemblies.

Since Andean potato weevil larvae complete their life cycle inside the potato, we obtained consent to collect adult weevil specimens that feed externally to avoid destroying farmers’ potato crops. The gut microbiota of these adult weevils resulted in much of the contamination observed using Kraken2 and BlobTools. Difficulty in processing permits for exporting these agricultural pest specimens led to lengthy storage in 95% ethanol at 4°C. This prolonged storage period led to DNA degradation making short-read sequencing the most viable option. A collection protocol designed for long-read sequencing, such as flash-freezing samples, would better facilitate future scaffolding and annotation of these four genome assemblies.

## Materials and Methods

### Sample Preparation and Sequencing

Weevil samples were collected from the wild in January 2018 across several regions of Peru: *P. vorax* (ID: 53D, Carhuaz, Lat. -9.26, Long. -77.59), *P. suturicallus* (ID: 28J, Jauja, Lat. - 11.71, Long. -75.52), *P. latithorax* (ID: 69J, Urubamba, Lat. -13.35, Long. -72.09), and *R. piercei* (ID: 90H, Puno, Lat. -15.68, Long. -70.14) and stored in 95% ethanol at 4°C. DNA was extracted from single adult weevils for each species using the EZNA Insect DNA Kit (Omega Biotek) and quantified using a Nanodrop 2000. Illumina paired-end libraries were prepared in the Roy J.

Carver Biotechnology Center at the University of Illinois at Urbana-Champaign with the Hyper Library Construction Kit (Kapa Biosystems) to obtain an average library fragment size of 500bp. Libraries were pooled and sequenced (2x150nt) on one lane of a S1 flowcell on the NovaSeq 6000 (Illumina, San Diego, CA). FastQ files were generated and demultiplexed with the bcl2fastq v2.20 Conversion Software (Illumina).

### Genome Assembly and Quality Assessment

Genome sizes were estimated for each species by counting 27-mers from the paired reads using Jellyfish v2.3.1 (68) and analyzing the histograms in GenomeScope 2.0 assuming a diploid sample (35). Raw reads from each sample were analyzed with Kraken2 v2.0.8 using the standard database and default parameters to identify possible contaminants (36). Reads classified as microbial were filtered from the *P. suturicallus* and *P. latithorax* sets prior to assembly. Raw reads for *R. piercei* and *P. vorax* were trimmed with fastp using default parameters with minimum length cutoff of 50nt and “--qualified_quality_phred 30” (69, 70).

Trimmed and/or filtered reads were assembled with MEGAHIT v1.2.9 using the option “-- presets meta-sensitive”, followed by filtering out contigs less than 1Kb for downstream work (71). Post-assembly contaminants were identified with BlobTools v1.1.1, which aggregates assembly metrics, contig read-depth, and taxonomic classification (72). Taxonomic classifications were determined using DIAMOND v.2.0.6 (73) blastx with the UniRef 100 protein database v.2020_06 (74). In weevils, contaminants could include pathogens such as bacteria or viruses, symbionts, and even food sources in the gut microbiota. Reads were aligned to each genome assembly with BWA v0.7.17 (75) using bwa mem default parameters, and a sorted and indexed bam file produced with Samtools v1.16.1 (76). Contigs identified as microbial in origin were filtered from the assembly, and any remaining duplicate contigs were identified and removed using the “Dedupe” feature from BBMap v38.36 with default settings (77).

The assembled contigs were then scaffolded in three sequential steps. First, the trimmed paired-end reads were combined with the MEGAHIT contigs in SOAPdenovo-fusion, a function of SOAPdenovo 2 v.r242, with insert size set to 500bp and K=49 (78). In the second scaffolding step, predicted coding regions from a *P. vorax* transcriptome served as guides for further scaffolding with L_RNA_Scaffolder (28, 79). RNASeq reads from *P. vorax* (NCBI SRA SRR8266148 and SRR8266149) were trimmed with fastp using the options “-- qualified_quality_phred 30 --average_qual 30 --trim_front1 10 --trim_front2 10 –trim_tail1 5 -- trim_tail2 5 --trim_poly_x –length_required 50” and then assembled with SOAPdenovo-Trans v1.03 with K=49 (80). The transcriptome consisted of 20,231 transcripts >1 kb summing to 40 Mb. Duplicated transcripts and non-beetle contaminant sequences were removed using methods as described above for the genome assemblies. Transcripts were then clustered with usearch v11.0.667_i86linux32 and options “-id 0.95 -centroids” (81). Open reading frames for each cluster were predicted with TransDecoder v5.5.0 with a minimum length of 100 (82). Each weevil genome assembly was aligned to the *P. vorax* set of ORFs using BLAT with default parameters to output a .psl file (83). The *P. vorax* .psl file and each genome assembly from SOAPdenovo-fusion were loaded into L_RNA_Scaffolder for scaffolding with default parameters.

For the final scaffolding step, RagTag v.1.1.1 was used for reference-guided genome scaffolding by testing the following candidate references: *Listronotus bonariensis* (Argentine stem weevil, Kuschel 1955, GCA_014170235.1), *Pissodes strobi* (White pine weevil, Peck 1817, GCA_016904865.1), *Rhynchophorus ferrugineus* (Red palm weevil, Olivier 1790, GCA_012979105.1), and *Sitophilus oryzae* (Rice weevil, Hustache 1930, GCF_002938485.1) (84). The Argentine stem weevil was selected as the reference for all RagTag-scaffolding as the BUSCO completeness scores were maximized with this species.

The programs BUSCO v5.2.2 (Benchmarking Universal Single-Copy Orthologs) with the odb Endopterygota lineage (85), QUASTv5.2.0 using default parameters (86), and the “stats” function of BBMapv.39.01 (87) were used as to assess assembly quality at each step. Genomic repetitive elements were first *de novo* modelled for each species with RepeatModeler2 (88) using -LTRStruct, followed by classification with RepeatMasker v4.1.0 using default parameters (89).

Genes involved with environmental receptors or digestive functions to sense or consume potato plants, as well as genes involved with insecticide resistance were retrieved from the Colorado potato beetle genome assembly Ldec_2.0 (NCBI: PRJNA854273) (34, 45). Using the more sensitive discontiguous MEGABLAST and other default parameters, the Colorado potato beetle genes were queried against the Andean potato weevil genome assemblies as this preset is intended for divergent species comparisons (90).

Whole-genome synteny analysis between each of the four Andean potato weevil species and *L. decemlineata* was performed with MUMmer4 (91) default parameters and visualized using CIRCOS default parameters (89). Due to the assembly limitations, discussed previously, complete genome annotations were not generated to avoid erroneous codon or gene placement (92). However, future long-read sequencing and improved scaffolding technologies paired with these current genome assemblies should make additional annotation achievable.

## Acknowledgements

We would like to thank the Peruvian potato farmers who graciously allowed us to work in their potato crops and collect these Andean potato weevil species. We are also thankful to Dr. Christopher J. Fields and Dr. Alvaro Gonzalo Hernandez at the Roy J. Carver Biotechnology Center (CBC) at University of Illinois Urbana-Champaign for their assistance with sequencing and facilitating the preliminary bioinformatics consultation for this project. This work was supported by the Wenner-Gren Foundation Dissertation Fieldwork Grant (Gr. 9907).

## Data Availability

The genomic data from this manuscript has been included as supplementary information or is openly available at: (NCBI: PRJNA1050318; *submission in review*). The assembled *P. vorax* transcriptome is openly available at: https://github.com/KelseyJorgensen/Premnotrypes_vorax_transcriptome.git.

## Supporting Information

**S1 Table. Genome assembly metrics with MEGAHIT**

**S2 Table. Genome assembly metrics post-decontamination**

**S3 Table. Genome assembly metrics with SOAPdenovo-fusion**

**S4 Table. Genome assembly metrics with L_RNA_Scaffolder**

**S5 Table. Comparison of references for use in RagTag scaffolding**

**S6 Table. Identified homologous genes with *L. decemlineata***

**S7 Table. Identified environmental receptors and insecticide resistance genes in *R. piercei* (90H)**

**S8 Table. Identified environmental receptors and insecticide resistance genes in *P. vorax (53D)***

**S9 Table. Identified environmental receptors and insecticide resistance genes in *P. latithorax* (69J)**

**S10 Table. Identified environmental receptors and insecticide resistance genes in *P. suturicallus* (28J)**

**S1 Figure. GenomeScope 2.0 predictions for Andean potato weevil assemblies.** GenomeScope outputs predicted sizes of A. 957 Mb for R. piercei (90H), B. 619 Mb for *P. vorax* (53D), C. 711 Mb for *P. latithorax* (69J), and D. 613 Mb for *P. suturicallus* (28J). Discordant features of the GenomeScope models were observed for all the outputs, particularly a lack of overlap between observed and modeled k-mers. This could imply one of several possibilities: that the genome size is much larger than expected, that the sequencing error rate is higher than expected, or that there is contaminating background DNA. The sequencing error rate was found to be negligible to absent for all four Andean potato weevil species. Three out of four weevil species did have larger final assemblies than initially expected, which likely greatly contributed to this GenomeScope discordance.

**S2 Figure. Identified contaminants with BlobTools.** BlobTools was used to identify any further contaminants within the mapped weevil assemblies after sensitive-MEGAHIT assembly. Mapped reads are shown by taxonomic group at the rank of ‘order’.

